# Examination of INPP5A in uveal melanoma uncovers novel calcium transients

**DOI:** 10.1101/2024.09.18.613756

**Authors:** Michael D. Onken, Kevin M. Kaltenbronn, Carol M. Makepeace, Kisha D. Piggott, Kendall J. Blumer

## Abstract

Uveal melanoma is a highly aggressive intraocular cancer that metastasizes in about half of patients whereupon it is inexorably fatal. Uveal melanomas (UM) are distinct from other melanomas because they are driven by constitutively activating mutation in the heterotrimeric G protein alpha subunits Gq (GNAQ) and G11 (GNA11). This results in constitutive production of inositol trisphosphate (IP3) by phospholipase C-beta downstream of Gq/11. In normal cells, increased IP3 causes calcium release from the endoplasmic reticulum, which would be cytotoxic if maintained chronically, but UM cells are able to survive constitutive IP3 production. INPP5A, which dephosphorylates and thus inactivates IP3, is highly upregulated in UM cells compared to other melanomas and another study has shown that INPP5A is necessary for UM cell survival. To understand the mechanism of calcium regulation in response to IP3, we collected single-cell calcium measurements and found that UM cells driven by constitutively active Gq/11 produce spontaneous calcium transients. These calcium oscillations are not seen in any other melanoma cell lines unless induced by an agonist, but they are present in patient UM tumor samples. Moreover, these calcium oscillations are lost in UM cells treated with the Gq/11 inhibitor FR900359, demonstrating their dependence on constitutive Gq/11 activity. We found that the INPP5A inhibitor YU144369 causes significant changes in calcium oscillations in UM cells, demonstrating a role for INPP5A in this system. INPP5A is tethered to membranes by C-terminal prenylation and palmitoylation, suggesting that localization may play a role in INPP5A regulation of IP3 levels. GFP-tagged INPP5A was localized to plasma membrane, nuclear envelop, endoplasmic reticulum, and lysosomes. Mutation of the palmitoylation site significantly reduced localization to the plasma membrane, while mutation of the prenylation site resulted in purely nucleoplasmic localization of INPP5A. These results demonstrate a role of palmitoylation in the regulation of INPP5A localization and mobilization in UM cells.

## Introduction

Uveal (ocular) melanoma (UM) is the most common cancer of the eye and the second most common form of melanoma after cutaneous (skin) melanoma (1). Almost half of patients diagnosed with a UM eye tumors go on to develop UM metastatic disease (2), which is fatal, and for which there are no curative therapies (3). Unlike cutaneous melanoma, metastatic UM does not respond to immune checkpoint blockade therapy, increasing the need for novel therapeutic approaches to treatment. Recent clinical trials using a bispecific T-cell activator (tebentafusp) have shown promise by extending survival in some patients but ultimately do not affect patient mortality from metastatic disease (4, 5).

Uveal melanomas arise from pigment cells in the choroid, ciliary bodies, or iris (collectively, the uvea). In 90% of UM patients, tumor initiation occurs when cells acquire oncogenic mutations in the G protein α-subunits Gαq (GNAQ) (6) or Gα11 (GNA11) (7). In another about 5% of patients, tumor initiation results from oncogenic mutations in the Gq/11-coupled leukotriene receptor (CYSLTR2) (8) or the immediate downstream effector of Gq/11, phospholipase Cβ (PLCβ, specifically PLCB4) (9). All of these mutations have the same downstream effect of constitutive activation of PLCβ, which breaks down phosphatidyl inositol bisphosphate (PIP2) to produce two important second messengers: diacylglycerol (DAG), and inositol trisphosphate (IP3). Several groups have demonstrated the importance of DAG in UM via its activation of protein kinase C and RASGRP3 (10) as they regulate MEK and ERK through the Ras-MAPK pathway in UM tumor oncogenesis and progression (11). However, clinical approaches with MEK and ERK inhibitors have been unsuccessful (12, 13).

A recent paper by Elbatsh, et al. (14) identified loss of the IP3 phosphatase INPP5A (inositol polyphosphate 5-phosphatase A) as synthetic lethal with constitutively active (ca) Gq/11 in UM cells (14). They show that knockdown or knockout of INPP5A from UM cell lines significantly reduces their survival, and this is specific to cell lines driven by caGq/11 (14). This prompted us to look more carefully at the IP3 side of the PLCβ equation. IP3 binds to IP3 receptors on the endoplasmic reticulum (ER) which function as calcium channels that release ER calcium stores into to cytosol on IP3 binding (15). INPP5A and the inositol pathway thus present new directions to explore for the identification of novel therapeutic targets for UM. We set out here to examine the function and regulation of INPP5A in UM cells driven by caGq/11.

## Results

### INPP5A is essential for survival and is highly expressed in UM cells

A recent study by Elbatsh, et al. demonstrated the dependence of Gq/11-driven UM tumors on expression of the gene inositol polyphosphate 5-phosphatase A (INPP5A) (14). We wanted to expand on these finding and determine the function and regulation of INPP5A protein in UM cells. We started with an unsupervised query for genetic dependencies of UM cell lines in the DepMap Portal (http://depmap.org), a comprehensive collection of high-throughput results from RNA interference and CRISPR gRNA libraries in a large, curated collection of cell lines representing several known cancers and diseases (16). DepMap identified INPP5A as the third most significant gene essential for UM cell survival (Figure 1A). This was particularly interesting because the other top gene hits in the DepMap analysis were directly involved either in Gq/11 signaling or melanocyte cell identity. INPP5A dephosphorylates IP3 produced by phospholipase C-beta (PLCβ) downstream of Gq/11 (17). In the context of constitutively active (ca) Gq/11, this dephosphorylation functions to reduce IP3 stimulation of IP3 receptor-calcium channels (IP3R) on the endoplasmic reticulum (ER) (Figure 1B). INPP5A dephosphorylating IP3 to IP2, which is then further degraded to IP1 (18), which is extremely high in MP41 (G11:Q209L) and MP46 (Gq:Q209L) caGq/11-driven cell lines compared to OCM-1A (BRAF:V600E) BRAF-driven cells (Figure 1C) and can be alleviated with the Gq/11 inhibitor FR900359 (FR) (19) (Figure 1C). We re-analyzed our published Mass-Spec data (11) to examine protein abundance of each of the three IP3R isoforms and for INPP5A in our cell lines. Protein levels for all three IP3R isoforms were significantly lower in caGq/11-driven UM cell lines (Figure 1D); whereas, INPP5A protein levels were significantly elevated compared to BRAF-driven OCM-1A cells (Figure 1E). We wanted to determine if our INPP5A findings in cell lines could be generalized to UM tumors. We turned to the Cancer Genome Atlas (TCGA; https://www.cancer.gov/tcga), which contains extensive genetic and clinical data on tumor samples from patients with UM (UVM; TCGA Pan-Cancer Atlas: 80 samples) and skin cutaneous melanoma (SKCM; TCGA Pan-Cancer Atlas: 448 samples). We found significantly higher INPP5A expression levels in UVM tumors compared to SKCM tumors (Figure 1F). We also analyzed mutation rates in both sets and found that the mutation rate for INPP5A was significantly lower (χ^2^ < 0.001) in UVM tumors. Moreover, the SKCM missense mutations were randomly scattered throughout the coding region, while the single reported missense mutation in UVM was in the C-terminal glutamine residue (Q412L) that would cause INPP5A to be geranylgeranylated instead of farnesylated (20). Because the geranylgeranyl moiety is more hydrophobic, this substitution would potentially stabilize association with membranes.

**Figure 1:**
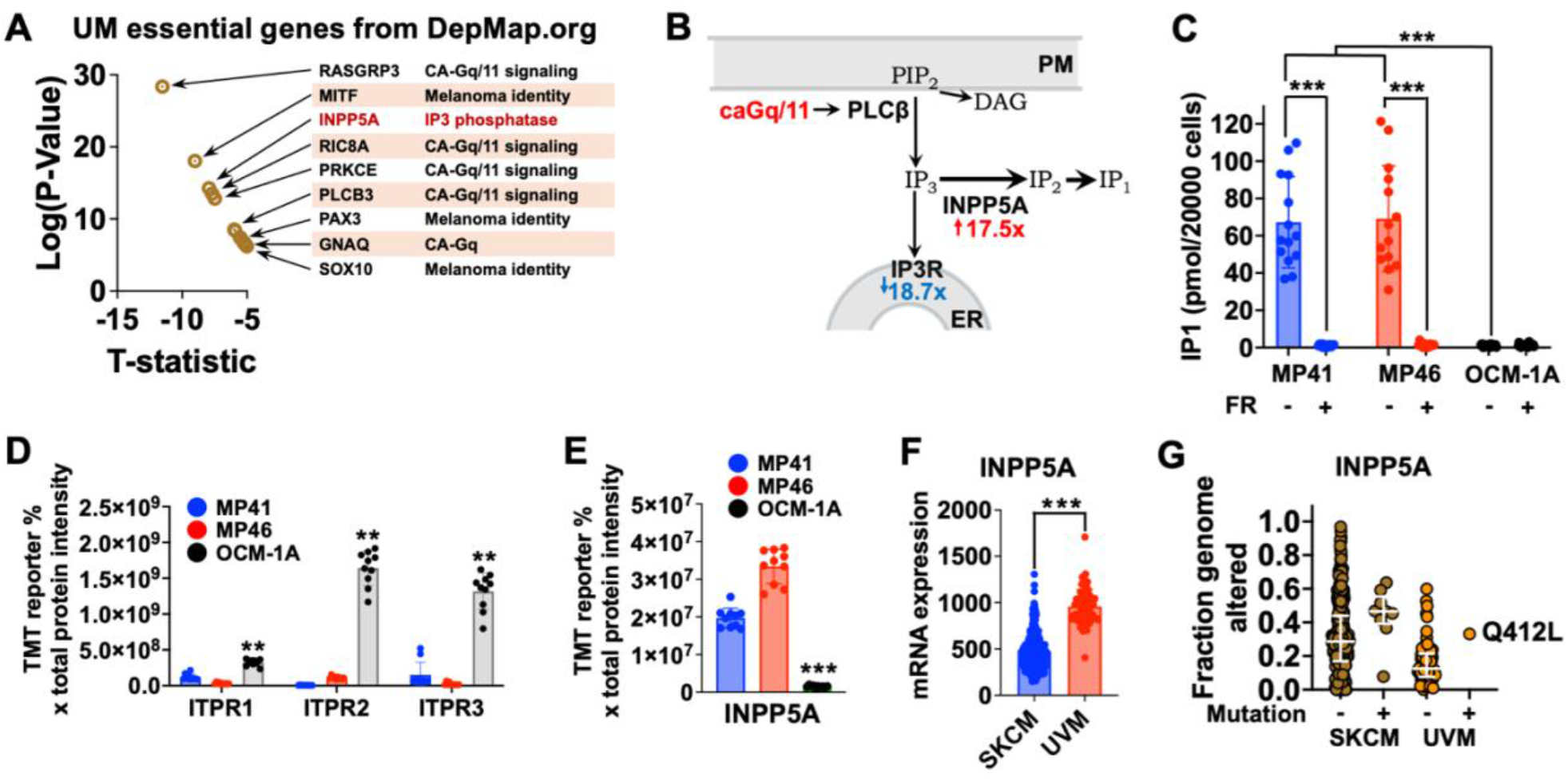
INPP5A is essential for survival and highly expressed in UM cells. **A)** The DepMap Portal (http://depmap.org) offers a comprehensive collection of RNA interference and CRISPR data for known genes in many cell lines representing several known cancers and diseases. We searched for genes that cause reduced survival when deleted from UM cell lines and graphed the top hits by significance and T-statistic. INPP5A was the third most significant hit and had a distinct function from the other top hits. **B)** Schematic of IP3 signaling downstream of constitutively active (ca) Gq/11. Phospholipase C-beta (PLCβ) is activated by caGq/11 to cleave phosphatidyl inositol bisphosphate (PIP2) in the plasma membrane (PM) into diacylglycerol (DAG) and inositol trisphosphate (IP3). IP3 binds to IP3 receptor-calcium channels (IP3R) on the endoplasmic reticulum (ER) to release ER calcium stores into the cytosol. INPP5A alleviates stimulation of calcium release by dephosphorylating IP3 to IP2, which is then further degraded to IP1. **C)** IP1 levels were quantified in UM cell lines treated with vehicle or FR (***: p<0.001). Protein abundance was calculated from our previously published Mass-Spec results for each of the three IP3R isoforms **(D)** and for INPP5A **(E)** in MP41 (G11:Q209L), MP46 (Gq:Q209L), and OCM-1A (BRAF:V600E) cells (**: p < 0.01; ***: p < 0.001). **E)** Gene expression data from the Cancer Genome Atlas (TCGA) were analyzed to compare INPP5A espression levels in skin cutaneous melanoma (SKCM) tumors to uveal melanoma (UVM) tumors (***: p < 0.001). **F)** TCGA were also analyzed to identify INPP5A mutations in human UM tumors. The mutation rate for INPP5A was remarkably low (1:80) with only a single reported missense mutation in the C-terminal glutamine residue (Q412L). The results shown here are in whole or part based upon data generated by the TCGA Research Network.

### Gq/11-driven UM cells exhibit spontaneous Ca^2+^ transients

The increased protein levels of INPP5A in caGq/11-driven UM cells should reduce cytosolic IP3 levels, while the decreased IP3R protein levels should reduce Ca^2+^ release from ER stores in response to IP3 (21). These adaptions to constitutively high PLCβ production of IP3 suggested a key role of calcium homeostasis in UM cell survival. To measure intracellular Ca^2+^ levels, we used Oregon green 488-BAPTA-1, AM (OGB1), a cell-permeable fluorescent dye that increases its fluorescent emission with increased Ca^2+^ binding. All cells showed similar baseline fluorescence after 1 hour incubation with OGB1. Surprisingly, both MP41 and MP46 UM cell lines exhibited spontaneous Ca^2+^ transients in a subset of individual cells (Figure 2A), which was not seen in OCM-1A cells (Figure 2A-C). Maxima for transient peaks varied among individual cells (Figure 2B), as did their periods of oscillation from transient to transient (Figure 2C). Overall mean heights and frequencies were similar between MP41 and MP46 cell lines, however the MP46 cell line showed a higher percentage of oscillating cells than MP41 (Figure 2E). Interestingly, MP41 and MP46 cells treated with the Gq/11 inhibitor FR lost their Ca^2+^ transients (Figure 2 D-E). Transients were also lost when cells were incubated in Ca^2+^-free medium for 1hour prior to imaging (Figure 2D).

**Figure 2:**
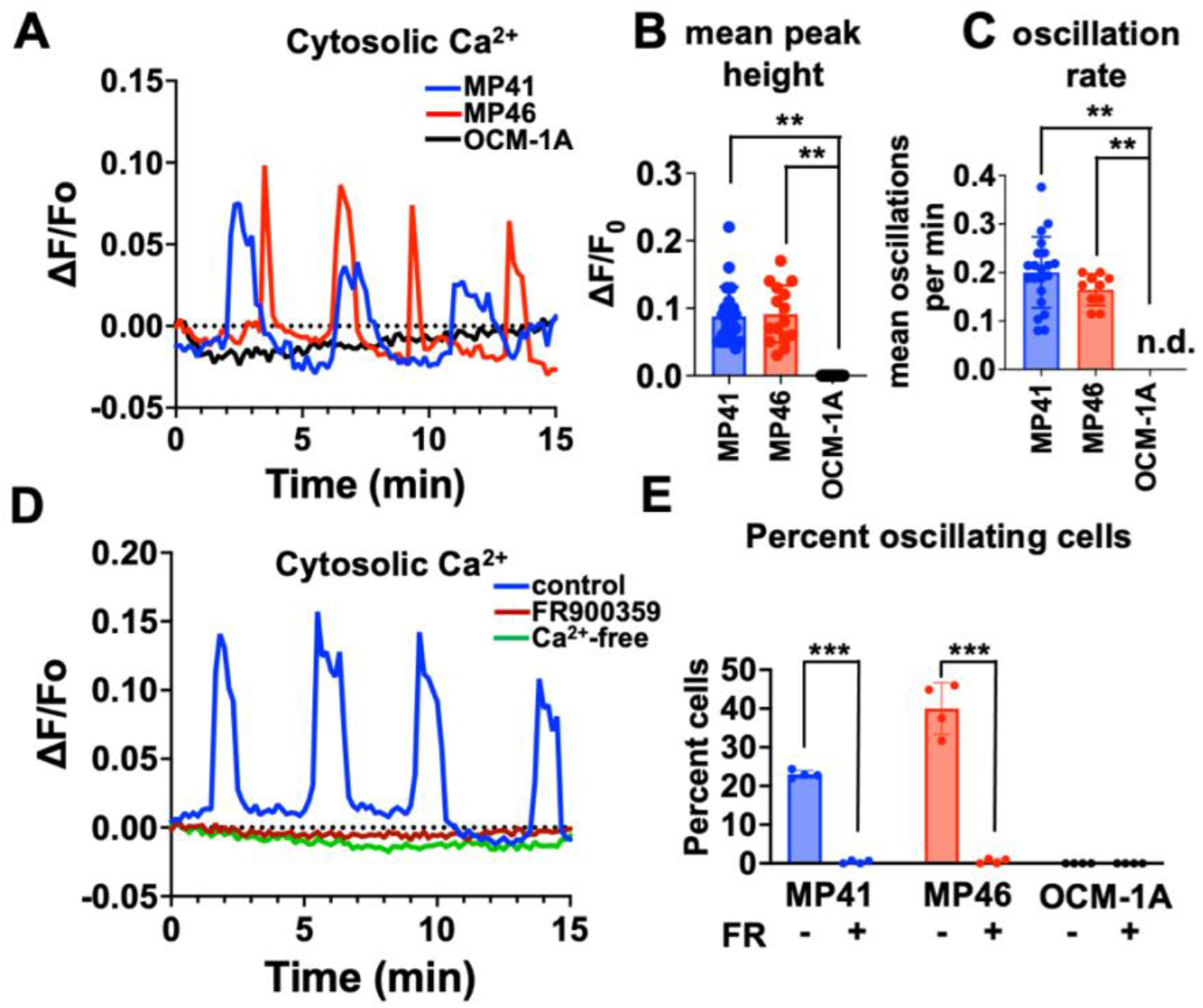
Gq/11-driven UM cells exhibit spontaneous Ca^2+^ transients. MP41 (G11:Q209L), MP46 (Gq:Q209L), and OCM-1A (BRAF:V600E) were stained with Oregon green 488-BAPTA and fluorescence emission associated with intracellular Ca^2+^ levels were measured over time. **A)** Representative Ca^2+^ traces for individual cells from each cell line. **B)** maxima for each transient peak were determined and mean peak heights were calculated for individual cells from each line. **C)** Periodicity was measured as the elapsed time from the start of one transient until the start of the subsequent transient for each individual cell trace and expressed as oscillations per minute. **D)** Representative Ca^2+^ traces for individual MP46 cells treated with vehicle or FR900359 for 24h, or placed in Ca^2+^-free medium for 1h. **E)** Fraction of cells per field that exhibited multiple transients were calculated for each cell line treated with vehicle or FR for 24h. For all panels, **: p < 0.01; ***: p < 0.001.

While the Ca^2+^ transients displayed by our two cell lines were provocative, we wanted to confirm that this behavior was present in real UM tumor cells. We collected a fine-needle aspiration biopsy sample from a primary human UM tumor, divided the sample to two imaging dishes, and maintained the cells in short-term culture to allow the tumor cells to adhere. The biopsy sample was treated with vehicle or FR for 24 hours and then incubated with OGB1 for Ca^2+^ imaging. Individual cells from the UM tumor biopsy showed spontaneous Ca^2+^ transients, similar to the MP41 and MP46 cell lines (Figure 3A). Interestingly, the biopsy samples displayed higher transient peaks than the cell lines (compare Figure 2B and Figure 3B; p < 0.01). The primary tumor samples also had a significantly higher proportion of oscillating cells than the cell lines (compare Figure 2E and Figure 3D; p < 0.01). Primary UM cells treated with FR stopped producing Ca^2+^ transients (Figure 3), similar to MP41 and MP46 cells. Although more samples are needed, these results suggest that Ca^2+^ transients are common among UM cells and are driven by caGq/11.

**Figure 3:**
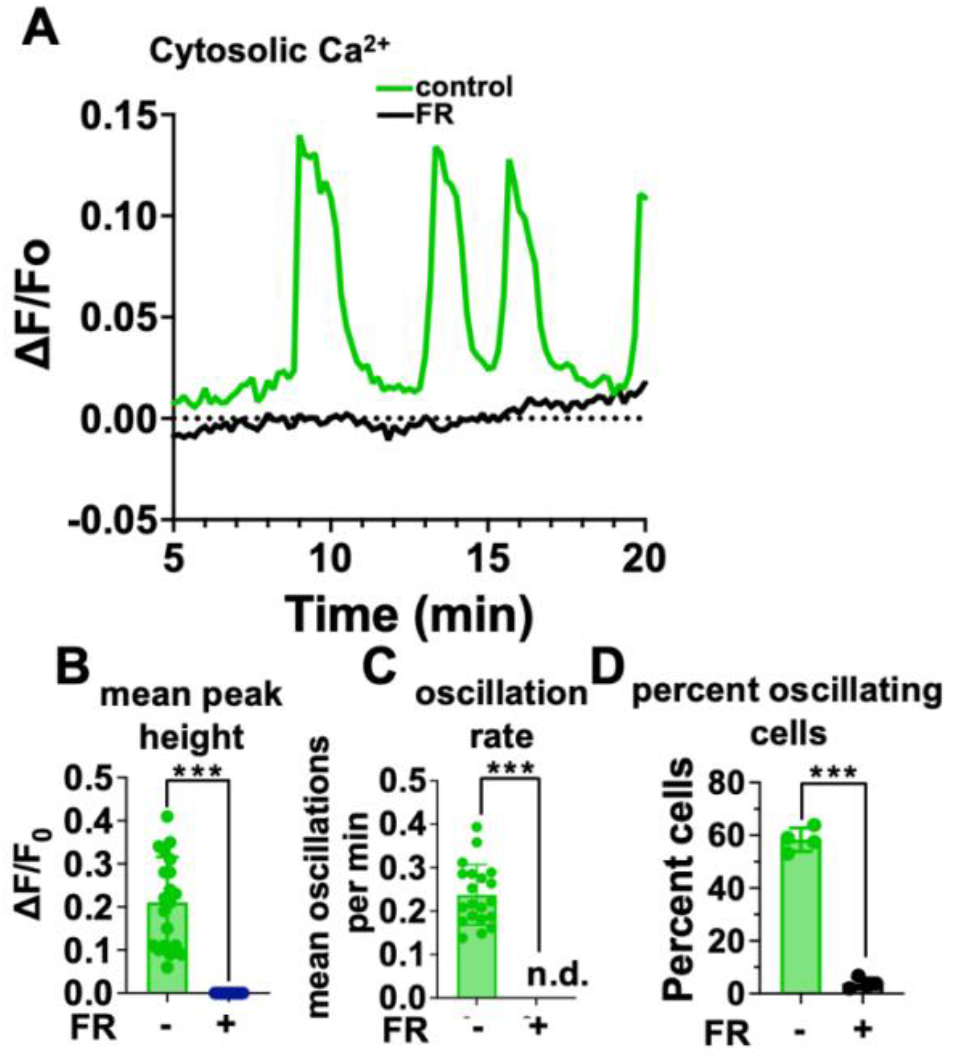
Primary human UM cells exhibit spontaneous Ca^2+^ transients. **A)** Representative Ca^2+^ traces for individual cells from a primary human tumor sample treated with vehicle or FR for 24 h. **B)** Mean peak heights were calculated for the primary tumor cells treated with vehicle or FR as in B. **C)** Periodicity was calculated for the primary tumor cells treated with vehicle or FR as in C. **D)** Fraction of cells per field that exhibited multiple transients were calculated for primary tumor cells treated with vehicle or FR. ***: p < 0.001.

### INPP5A regulates Ca^2+^ transient behavior in UM cells

INPP5A regulates IP3 levels through dephosphorylation, which could also affect Ca^2+^ transient behavior in UM cells. MP46 cells were assayed with OGB1 as previously, but dishes were injected with vehicle or the inositol polyphosphate 5-phosphatase inhibitor YU144369 (YU) (22) at 1 min after start time (Figure 4A, arrow). Addition of YU resulted in an immediate burst of cytosolic Ca^2+^ in all MP46 cells (Figure 4A, highlighted in purple). After this burst, individual cells continued to exhibit Ca^2+^ transients (Figure 4A), however the post-burst Ca^2+^ transients were shorter (Figure 4B) and more frequent (Figure 4C) than control cells – the initial burst was not included in either the peak height or frequency calculations. Further, treatment with YU resulted in almost all of the cells in each field to exhibit transient behavior after the initial burst. These results demonstrate that IP3 regulation by INPP5A contributes to caGq/11-driven Ca^2+^ transient behavior in UM cells.

**Figure 4:**
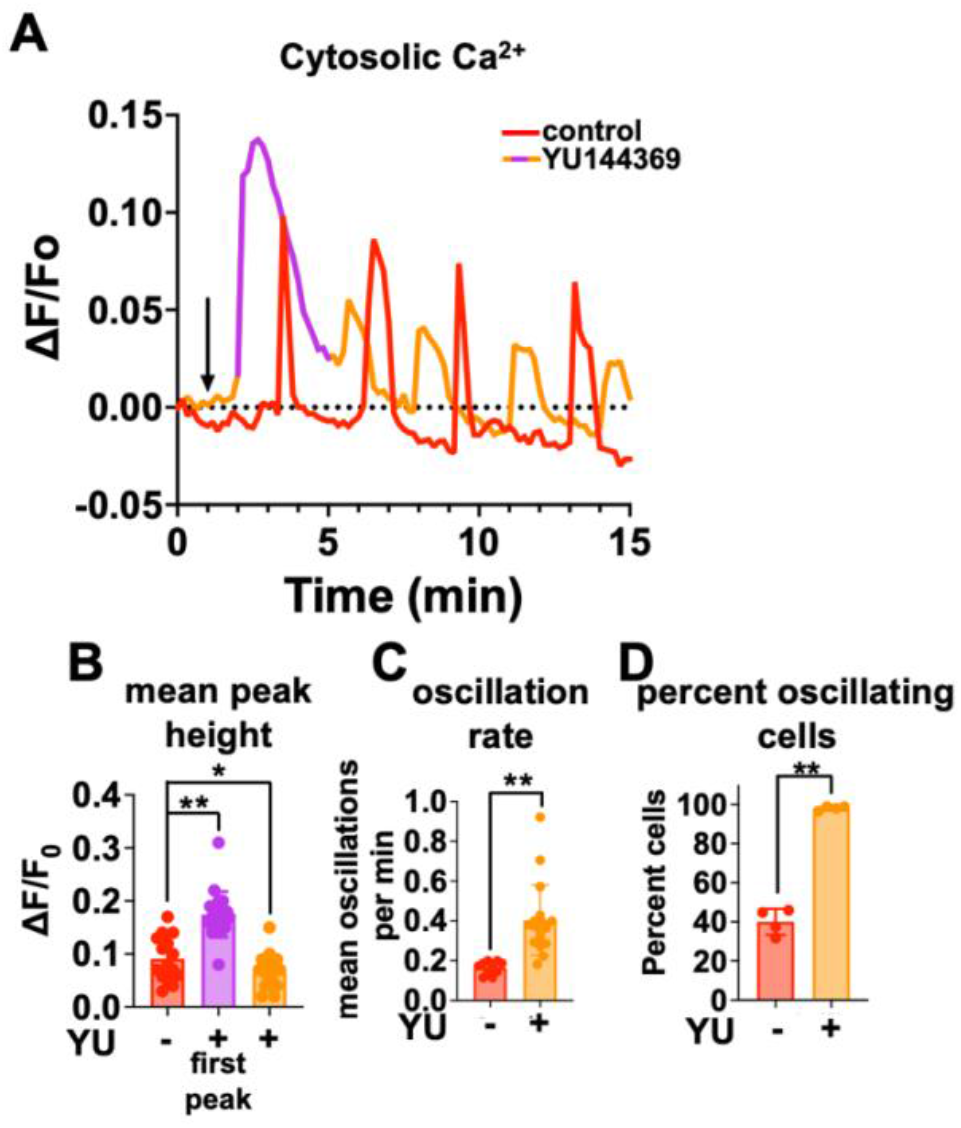
Inhibition of INPP5A acutely affects Ca^2+^ transients. **A)** Representative Ca^2+^ traces for individual MP46 cells injected with vehicle or YU144369 at 1 min after start time (arrow). **B)** An initial Ca^2+^ burst was seen in all cells immediately after YU injection and this was determined separately as the first peak. Mean peak heights for the subsequent transients were calculated as in B. **C)** Periodicity was measured as in C, but the initial burst was not included in the calculations for the YU-treated cells. **D)** Fraction of cells per field that exhibited multiple transients were calculated for vehicle-treated cells and YU-treated cells after the initial Ca^2+^ burst. For all panels, *: p < 0.05; **: p < 0.01.

### INPP5A localizes to cell membranes

Prior studies have shown lipidation of the C-terminus of INPP5A (20). This would suggest interaction with and localization to membranes when expressed in cells. We constructed lentiviral vectors expressing GFP-INPP5A fusion proteins and infected HEK293 cells and performed live-cell imaging on a spinning-disk confocal fluorescent microscope to determine the subcellular localization of INPP5A in human cells. Exogenously expressed GFP-INPP5A localized mostly to the plasma membrane and nuclear envelope (Figure 5A, arrows) as well as to distinct cytoplasmic structures rather than being diffusely cytosolic. To identify the cytoplasmic structures labeled by GFP-INPP5A, we first co-expressed an ER-specific fusion construct mCherry-SEC61B (23). Sec61β is the beta subunit of the translocon protein complex, which is an integral trans-membrane component of the ER. Co-expression of GFP-INPP5A and mCherry-SEC61B shows clear overlap of the two fluorescent signals on ER (Figure 5B), however GFP-INPP5A also showed localization to several puncta and larger vesicles that were not labeled by mCherry-SEC61B. The larger vesicular structures looked like lysosomes, so we co-expressed GFP-INPP5A with a Lysosomal-associated membrane protein 1-RFP (LAMP1-RFP) fusion protein construct (24). LAMP1 is an integral membrane protein localized to vesicles in the lysosomal pathway. Co-expression of GFP-INPP5A and LAMP1-RFP identified the INPP5A-positive puncta and vesicles to be components of the lysosomal pathway (Figure 5C). These results demonstrate the localization of INPP5A to intracellular membranes, although by overexpressing INPP5A may alter the relative contributions of each compartment to endogenous INPP5A pools.

**Figure 5:**
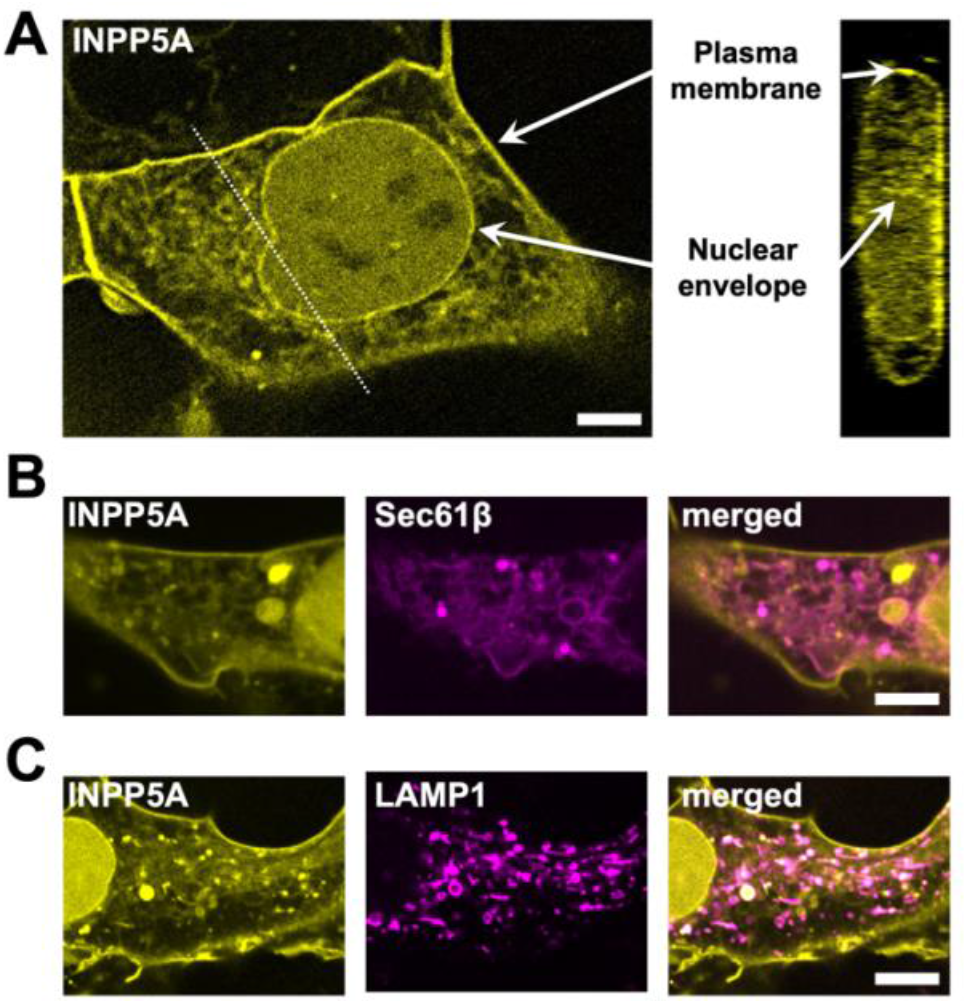
INPP5A localizes to cell membranes. HEK293 cells expressing GFP-INPP5A were imaged on a spinning-disk confocal fluorescent microscope. **A)** Exogenously expressed GFP-INPP5A localizes to the plasma membrane and nuclear envelope (arrow) and distinct cytoplasmic structures. Scale Bar: 5 µm. **B)** Sec61β is the beta subunit of the translocon protein complex, which is an integral trans-membrane component of the ER. Co-expression of GFP-INPP5A (left panel) and mCherry-SEC61B (middle panel) shows clear overlap of the two fluorescent signals on ER (right panel). Scale Bar: 5 µm. **C)** Lysosomal-associated membrane protein 1 (LAMP-1) is an integral membrane protein localized to vesicles in the lysosomal pathway. Co-expression of GFP-INPP5A (left panel) and RFP-LAMP1 (middle panel) identifies INPP5A-positive puncta and vesicles targeted to the lysosomal pathway (right panel). Scale Bar: 5 µm.

### Localization of INPP5A to membranes is regulated by prenylation and palmitoylation

The C-terminal domain of INPP5A ends in a CAAX box (Figure 6A), which signals terminal prenylation (20), preceded by a Cysteine residue, which becomes a target of palmitoylation after C-terminal prenylation (20). Previous studies have shown that the terminal glutamine at residue 412 directs C-terminal farnesylation rather than geranylgeranylation of INPP5A (20). We measured the extent of palmitoylation of INPP5A by using 17-octadecynoic acid (17-ODYA), a functionalized homolog of palmitic acid (25). After incorporation, 17-ODYA can be labeled with TAMRA dye to identify proteins that have been palmitoylated (25). We used HEK293 cells expressing wildtype FLAG-INPP5A (WT) or mutants lacking the palmitoylation (C408S) or farnesylation (C409S) sites and treated them with vehicle or the depalmitoylase inhibitor hexadecylfluorophosphonate (HDFP) (26). By first immunoprecipitating lysates with anti-FLAG antibodies, the TAMRA signal was restricted to FLAG-tagged proteins (Figure 6B). Western blotting with an antibody against INPP5A confirmed showed overlap between the TAMRA signal and INPP5A protein (Figure 6B). HDFP increased the steady state level of 17-ODYA incorporation about 2-fold by fluorescence densitometry. Palmitoylation is a highly regulated process with constant turnover by depalmitoylases and palmitoylases. Palmitate turnover on INPP5A was measured by 17ODYA pulse-chase (Figure 6C) and found to have a half-life around 71 min. (range: 58 min. to 88 min. as calculated by single-phase decay regression analysis). The turnover of palmitoylation of INPP5A suggests a regulatory role in targeting and INPP5A to specific compartments. To test this, we used expressed GFP-INPP5A(WT) or mutants lacking the palmitoylation (C408S) or farnesylation (C409S) sites in HEK293 cells and performed live-cell imaging on a spinning-disk confocal fluorescent microscope as before. Sub-cellular localization of palmitoylation and farnesylation site mutants showed significant differences with wildtype INPP5A (Figure 6D). Mutation of the palmitoylation site resulted in reduced plasma membrane localization, reduced ER localization, and increased localization to the nuclear envelope. By contrast, mutation of the farnesylation site eliminated all cytoplasmic INPP5A and instead caused the accumulation of INPP5A in the nucleoplasm. These results demonstrate that both palmitoylation and farnesylation are essential for the proper targeting if INPP5A to its normal sub-cellular compartments.

**Figure 6:**
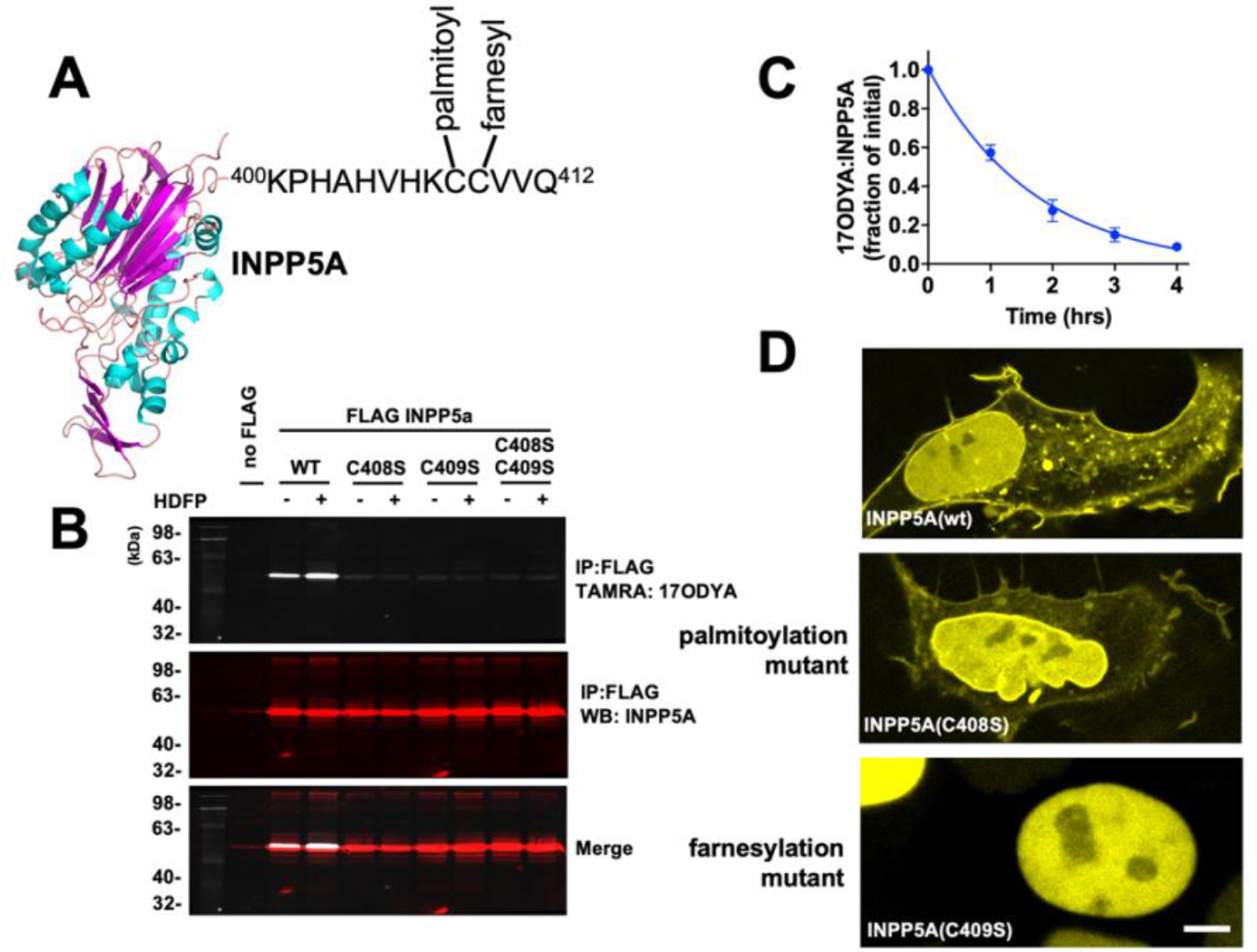
Localization of INPP5A to membranes is regulated by prenylation and palmitoylation. **A)** The C-terminal domain of INPP5A (AlphaFold prediction shown) ends in a CAAX box, which signals terminal prenylation, preceded by a Cysteine, which becomes a target of palmitoylation after C-terminal prenylation. Previous studies have shown that the terminal glutamine at residue 412 directs C-terminal farnesylation rather than geranylgeranylation of INPP5A. **B)** Direct measurement of palmitoylation of INPP5A was carried out using 17-octadecynoic acid (17-ODYA) followed by TAMRA labeling in HEK293 cells expressing wildtype FLAG-INPP5A (WT) or mutants lacking the palmitoylation (C408S) or farnesylation (C409S) sites. Immunoprecipitation with anti-FLAG antibodies was used to improve specificity for the transgene. **C)** Palmitate turnover on INPP5A was measured by 17ODYA pulse-chase. **D)** Sub-cellular localization of palmitoylation and farnesylation site mutants of GFP-INPP5A were imaged in HEK293 cells.

## Discussion

### UM cells exhibit a common adaptive mechanism to survive chronic PLCβ activation

Data from the DepMap project, TCGA, and our own expression data agree with the findings of Elbatsh, et al. (14): INPP5A is significantly upregulated in UM tumors compared to cutaneous melanomas; INPP5A is rarely mutated in UM, suggesting selective pressure to keep the gene intact; and loss of INPP5A is lethal to UM cells driven by caGq/11. Interestingly, we also saw significant reduction in all three IP3R isoforms at the protein level compared to cutaneous melanomas. This is particularly interesting because mRNA levels of the IP3Rs were not significantly different between UVM and SKCM tumors in the TCGA data. This would suggest that the adaptive reduction in IP3R protein is post-translational, likely through the Erlin complex that regulates IP3R ubiquitination and proteosomal degradation (27). Further research into this phenomenon may be merited as another avenue for therapeutic targeting of UM.

### UM cells exhibit unique spontaneous calcium transients

Our preliminary attempts at examining intracellular calcium changes in UM cells using a plate reader to measure bulk calcium levels failed multiple times, instead yielding noisy, uninterpretable data. After long discussions, we switched to single-cell calcium measurements under fluorescent microscopy instead of bulk well-based assays. We found that the noisy results were due to the asynchronous nature of the calcium transients displayed by a portion of the cells in each well. This single-cell approach completely changed our perception of calcium activity in UM cells and has given us the tools to dive deeper into how calcium transients are regulated in UM cells. From a genetic perspective, it is clear that constitutively active Gq/11 drives these calcium transients in UM cells, because they are not present in cells not driven by caGq/11, and because they go away with inhibition of Gq/11 by FR. This may not be so surprising in the broader context of Gq/11 activation: many other cells, especially in the neural lineages, demonstrate similar calcium transients in response to ligands that activate GPCRs that signal through Gq/11 (28). It is possible that the transients we see in UM cells represent the normal cellular responses to Gq/11 activation and may reveal an evolutionary mechanism to present cytosolic calcium toxicity and to maintain ER calcium stores. We continue to expand our assays to more cell types and other inhibitors in the signaling pathway, especially in light of the strong effects of INPP5A inhibition on UM calcium transients.

### INPP5A is directed to intracellular membranes by palmitoylation and farnesylation

The specific lipidation of the C-terminus of INPP5A has been shown previously (20). We now show the specific compartmental localization of INPP5A and how those lipid modifications regulate targeting INPP5A to specific compartments. One major pitfall of our approach is the use of overexpression constructs, which may not fully represent the sub-cellular localization on endogenous INPP5A. For example, the localization of overexpressed INPP5A to endosome and lysosomes may be due more to its overexpression on the plasma membrane than to specific targeting (29). Ultimately, the overexpression experiments illuminate where INPP5A can localize in the cell and not necessarily the levels of endogenous protein on each compartment. Most striking though are the changes in localization with mutation of the palmitoylation and farnesylation sites. The loss of cytoplasmic INPP5A(C409S) demonstrates the requirement of prenylation of INPP5A for its cellular function. Moreover, the C409S prenylation mutation also prevents palmitoylation, demonstrating that INPP5A needs the initial lipid anchor intact for membrane-bound palmitoyl transferases to recognize INPP5A and further lipidate it (25).

## Methods

### Cell lines and reagents

FR900359 was purified from A. crenata according to published methods (30). The human UM PDX-derived cell lines MP41 (ATCC Cat# CRL-3297, RRID:CVCL_4D12) and MP46 (ATCC Cat# CRL-3298, RRID:CVCL_4D13) (31) were purchased from ATCC (Manassas, VA). The human UM cell line OCM-1A (RRID:CVCL_6934) was derived by and the generous gift of Dr. June Kan-Mitchell (Biological Sciences, University of Texas at El Paso). All cell lines were grown at 37°C in 5% CO2 in RPMI 1640 medium (Life Technologies, Carlsbad, CA) supplemented with antibiotics and 25% FBS (MP41 and MP46) or 10% FBS (OCM-1A). Cells were not grown above passage 35. GFP-INPP5A was inserted into the pCDH-CMV-MCS-EF1-Puro lentivector expression plasmid, and lentiviral particles were produced as described previously (31). Cells were incubated with lentivirus in growth medium containing 10 µg / mL protamine sulfate for 24 hours before changing to fresh growth medium. At three days post infection, cells were moved to growth medium containing 1.5 µg / mL puromycin to select for stably integrated cells. TransIT-LT2 transfection reagent (Mirus Bio, Madison, WI) was used for all transient transfection according to manufacturers’ protocol. LAMP1-RFP (Addgene#1817) and mCherry-SEC61B (Addgene #90994) expression constructs were the generous gifts of Dr. David Kast.

### Patients and sample collection

Human tissue samples were obtained with patient written informed consent and with approval of the institutional review board of Washington University in St. Louis. Fine-needle aspiration biopsies of primary UM tumors were performed as part of standard of care, which also collected biopsies for cytological evaluation of tumor cells and molecular classification (Castle Biosciences). All samples were collected directly into 2 mL growth medium in the operating room before being transported to the laboratory. Biopsy samples were centrifuged and resuspended in 500 µL growth medium and then equally divided onto fibronectin-coated MatTek glass-bottom dishes (P35G-1.5-14-C) and grown overnight in a 5% CO2 TC incubator. Growth medium for primary UM samples is MDMF medium which consists of HAM’s F12 (Lonza, Walkersville MD, USA) supplemented with 1 mg/mL BSA (Sigma-Aldrich, St Louis MO, USA), 2 mM L-glutamine (Lonza), 1X SITE (Sigma-Aldrich), 1x B27 (Gibco, Carlsbad CA, USA), 20 ng/mL bFGF (PeproTech Inc, Rocky Hill NJ, USA), and 50 μg/mL Gentamicin (Sigma-Aldrich) (ref). Cells were allowed to attach to the substrate before 2 mL of fresh medium was added to each dish.

### IP1 assays

Accumulation of IP1 in UM cells was measured using the IP-One kit (CiSbio, Inc; catalog number 62IPAPEB) according to the supplier’s instructions. 20,000 MP41 cells, 20,000 MP46 cells and 20,000 OCM-1A cells were seeded into white-bottom tissue culture grade 384-well plates. Following an overnight incubation, cells were treated with FR or vehicle (DMSO) and returned to the incubator. The next day, stimulation buffer was added for 1 hour, after which IP1-d2 and Ab-Cryp were added, and the cells were incubated at room temperature for 60 min. Plates were read in a Synergy H4 Hybred Reader (BioTek, Winooski, VT, USA). Standard curves were generated using reagents supplied with the kit.

### Intracellular Calcium assays

Cells were plated on MatTek dishes with coverglass bottoms coated with fibronectin and grown overnight at 37°C. Cells were rinsed in DPBS at RT and then loaded with Oregon green 488-BAPTA-1, AM in FluoroBrite DMEM containing pluronic F-127 detergent for 1 hr at RT. Staining solution was removed and cells were rinsed twice with DPBS at RT to remove extracellular stain and then placed in FluoroBrite DMEM for imaging at RT. Time-lapse images were collected at 10 second intervals on an Olympus IX83 inverted microscope with a Lumencor Sola light engine and a 475nm excitation / 525nm emission filter set using a 10X objective and a Hamamatsu ORCA-FLASH4.0 sCMOS camera. All images were collected at the full camera dynamic range and the same thresholds were used to process all images. A 25-pixel radius circle was used to collect fluorescent intensity data from each cell at each time point, which was then converted to change in fluorescence over initial fluorescence (ΔF/F_0_).

### Fluorescence Imaging

Cells were plated on MatTek dishes with coverglass bottoms coated with fibronectin and allowed to adhere overnight at 37°C. Imaging was performed on a Nikon Ti2 inverted microscope equipped with a 60X oil immersion objective and a Yokogawa CSU-W1 spinning disk confocal attached to a Hamamatsu ORCA-FLASH4.0 sCMOS camera. Image stacks were captured at 16-bit 2048 × 2044 resolution with an axial spacing of 0.5 µm using the Nikon Elements Software package. All images were collected at the full camera dynamic range and the same thresholds were used to process all images. Specific Z-slices were chosen for figure presentation that represented the z-midline of the cell body. Orthogonal reslicing was performed in ImageJ.

### Palmitate assays and immunoprecipitation

HEK293 cells stably expressing FLAG-INPP5A were labeled with 25 μm 17-ODYA for 6 h in DMEM (Invitrogen) supplemented with 5% FBS, 0.1 mm sodium pyruvate, and 0.1 mm nonessential amino acids in the presence or absence of 20 µM HDFP. For pulse-chase experiments, 17-ODYA-treated cells were washed with phosphate-buffered saline (PBS) prior to being chased with media containing 200 μm palmitate for the indicated periods of time. At the end of each experiment, cells were washed and suspended in lysis buffer (PBS supplemented with 2.5 μm PMSF, 1× EDTA-free Complete Protease Inhibitors, and 1% Triton X-100). Cleared lysates were immunoprecipitated overnight at 4 °C with mouse anti-FLAG M2-agarose for 2 h at 4 °C. After three washes with lysis buffer, beads were suspended in PBS. Click chemistry reaction protocols were adopted from previous publications (30, 31). Immunoprecipitated samples were reacted with click chemistry reagents (20 μm TAMRA-azide, 1 mm tris(2-carboxyethyl)phosphine hydrochloride, 100 μm tris-(benzyltriazolylmethyl)amine, and 1 mm CuSO4) for 1 h at room temperature with periodic mixing. Reactions were stopped by addition of reducing SDS-PAGE sample buffer and boiling for 5 min at 100 °C. Samples were separated by SDS-PAGE and analyzed by in-gel fluorescence analysis (Typhoon 9400 laser scanner, GE Healthcare) and Western blotting. Bands of interest were quantified using ImageJ for TAMRA fluorescence. Western blot analyses were performed according to the following protocol. Samples were separated by SDS-PAGE, transferred to PVDF (Millipore, catalog number IPVH00010), blocked for at least 1 h using 5% (w/v) milk in TBST (25 mm Tris (pH 7.2), 150 mm NaCl, 2.7 mm KCl, 0.1% (v/v) Tween 20), incubated with mouse anti-FLAG primary monoclonal antibody in blocking buffer for at least 2 h, washed three times with TBST, incubated for 1 h with appropriate HRP-conjugated secondary antibodies diluted in blocking buffer, and washed three times with TBST. Membranes were then incubated with IRDye 800 Goat anti-mouse (LI-COR, Lincoln, NE). Following incubation, membranes were washed at least three times with TBST and signals were detected using LI-COR Odyssey model 9120 imaging system (LI-COR). All quantified Western blot signals were within the linear range of the detection system as determined by an independent standard curve.

### Data analysis and statistical analysis

Data from the Cancer Dependency Map Project were collected through the DepMap Portal (https://depmap.org/portal/). Data downloaded from DepMap were graphed in GraphPad Prism (version 10.3.1). Protein abundance data for MP41, MP46, and OCM-1A cell lines were calculated previously (ref) and re-graphed in GraphPad Prism. Data from the Cancer Genome Atlas Project (TCGA; https://www.cancer.gov/tcga) were collected using the cBioPortal Cancer Genomics webtool (https://www.cbioportal.org) using a combined set of the UVM (TCGA Pan-Cancer Atlas: 80 samples) and SKCM (TCGA Pan-Cancer Atlas: 448 samples) samples and searching by Gene (INPP5A). The data were downloaded as text files and re-graphed in GraphPad Prism. All statistical analyses were performed in GraphPad Prism (version 10.3.1). Average data values and error bars on graphs represent means with standard deviation, except for Figures 1F and 1G (TCGA data) which represent median and interquartiles. Stars indicate significance as determined by statistical analysis (*<0.05, **<0.01, ***<0.01). Data for treated and untreated samples were analyzed via unpaired t-tests or Mann-Whitney tests. Non-linear regression one-phase exponential decay was used for palmitoylation pulse-chase half life estimation.

## Supporting information

MP46 cells vehicle treated

MP46 cells YU144369 treated

